# Intracellular interaction of FNDC1/FN1/AR/cMYC in prostate cancer cells

**DOI:** 10.1101/2023.02.23.529660

**Authors:** Priyanka Ghosh, Olorunseun O. Ogunwobi

## Abstract

MicroRNAs play an important role in modulating normal cellular functions through protein-protein interaction in addition to regulating gene expression via mRNA degradation or translational repression. *PVT1* is a non-protein coding gene that encodes six annotated microRNAs (miRNAs), including miR-1207-3p, a demonstrated modulator in PCa. MiR-1207-3p is significantly underexpressed in PCa cells when compared to normal prostatic cells and directly targets fibronectin type III domain containing 1 (FNDC1), which was found to be overexpressed in PCa cells along with concomitant overexpression of fibronectin (FN1)/androgen receptor (AR)/c-MYC. To better understand the role of FNDC1/FN1/AR/c-MYC in PCa progression we examined the interaction between FNDC1/FN1/AR/c-MYC. Coimmunoprecipitation study showed direct physical interaction between FNDC1/FN1/AR/c-MYC in PCa cells and FN1/AR/cMYC in normal prostatic cell. Knockdown of FN1, AR and cMYC not only confirmed FN1/FNDC1/AR/cMYC interaction but also gave rise to the possibility of a complex. We also examined the spatial localization of the FNDC1/FN1/AR/c-MYC pathway by performing immunofluorescence staining in PCa cells. Single staining analysis revealed that FNDC1, AR and c-MYC localize to the nucleus and cytoplasm while FN1 localizes only to the cytoplasm in PCa cells but not in the non-tumorigenic prostate cells. However, our findings show that miR-1207-3p has no direct effect on the FNDC1/FN1/AR/cMYC interaction. Understanding this molecular interaction can reveal additional insights into the role of FNDC1/FN1/AR/cMYC interaction in PCa progression.

## Introduction

The American Cancer Society estimates 268,490 cases of prostate cancer (PCa), the second most common cancer among the men of America, in 2022 [49]. *PVT1* located at 8q24 is overexpressed and regulates tumor progression in prostate cancer. In addition to a long non-coding RNA, PVT1 also encodes six miRNAs including miR-1207-3p. MiR-1207-3p a tumor suppressor targets FNDC1, which is overexpressed in prostate cancer cells. Moreover, miR-1207-3p in PCa cells regulates proliferation, migration and apoptosis via direct molecular targeting of fibronectin type III domain containing 1 (FNDC1) and consequent loss of expression of fibronectin (FN1), androgen receptor (AR) and cMYC [30].

FNDC1 is known to over express in a multitude of cancer, such as gastric cancer, breast cancer and colorectal cancer etc including prostate cancer. [53, 54, 55]. It is known to regulate proliferation, migration and apoptosis in cancer cells by interacting with signaling molecule important in cancer progression. This pathway includes the Wnt signaling pathway and PI3K/Akt.

Fibronectin, an important part of the extracellular matrix belongs to the family of glycoproteins. Two identical subunits of 220 KDa form the high molecular weight (440KDa) FN1 protein which is covalently bound at the CTD with di-sulfide bonds. The multi-modular structure of FN1 is composed of type I, type II and type III repeats. These repeats further form the N-terminal domain (NTD: FNI_1-9_), central binding domain (CBD: FNIII_1-12_) and the heparin binding domain (HEP-II: FNIII_12-14_). Human cellular fibronectin has 20 different isoforms which is generated via the alternative splicing of the EIIIA, EIIIB and EIIICS domains during transcription [47].

AR a nuclear receptor belongs to the superfamily of thyroid, estrogen, progesterone, glucocorticoid and mineralocorticoid receptors [37]. Its overexpression is common in 30%-50% of CRPC patients which is a result of the AR gene amplification [51, 52]. Androgen receptors constitute the N terminal domain (NTD), DNA binding domain (DBD), ligand binding domain (LBD) and C terminal domain (CTD). The NTD plays an important role as a promoter in ligand independent AR activity. It also aids in nuclear localization, binds co-regulatory interactors and is necessary for tumorigenic proliferation [38,39,40, 41,42]. The various AR splicing variants observed in nature have their CTD or LBD truncated [46, 47, 48, 49]. Apart from the 20 or more AR isoforms known there are two more that is a result of the NTD domain truncation. These are the AR-B which is the full length and the AR-A which lacks the first 187 amino acid in the N terminal domain [35]. The AR-A is 2.38-2.81 and the AR-B 1.37-1.47 times more expressed in prostate cancer cells compared to normal and benign hyperplasia cells [36].

cMYC a member of the basic-helix-loop-helix-leucine-zipper family of transcription factors is aberrantly overexpressed in over 70% human cancers [45]. In addition to the N-terminal that acts as the transcription regulatory domain and C-terminal domains that take part in DNA binding, cMYC is composed of a central domain that contains PEST degradation and nuclear localization signal. The two commonly known cMYC isoforms are isoform1 that is short and initiates from AUG and the long isoform2 that initiates from CUG [43, 44]. However, a newer study has shown a third isoform of cMYC term cMYC-S. These isoforms mainly differ in their N-terminal region and differs functionally in deciding apoptosis [46].

## Materials and Methods

### Cell Culture

The LNCaP cell line modelling metastatic prostate cancer, was purchased from the American Type Culture Collection and cultured in RPMI 1640 supplemented with 10% heat inactivated FBS, and 1% penicillin/streptomycin. The C4-2B cell line is a well validated and widely recognized model of CRPC obtained from MD Anderson Cancer Center under a material transfer agreement with Hunter College of The City University of New York. C42B was maintained in DMEM combined with Ham’s F12 in a 3:2 ratio, supplemented with 10% heat-inactivated FBS, 1% penicillin/streptomycin, insulin (5µg/ml), triiodothyronine (13.65 pg/ml), human apo-transferrin (4.4 µg/ml), d-biotin (0.244 µg/ml), and adenin (12.5 µg/ml). The non-tumorigenic prostate cell, RWPE1 was cultured in Keratinocyte SFM Medium supplemented with BPE, EGF and 1% penicillin/streptomycin. All cells were maintained at 37°C in a humidified atmosphere with 10% CO_2_.

### Chemical Crosslinking and Coimmunoprecipitation

In order to crosslink FN1 a transmembrane protein to its interactors, cells were fixed with the cell impermeable bissulfosuccinimidyl suberate (BS3) crosslinker at 5 mM (21580, Thermo Scientific, USA) for half hour at room temperature followed by quenching with 20 mM Tris-HCl (pH 7.5) for 15 minutes. Protein G Dynabeads® co-immunoprecipitation kit (Invitrogen, UK) was used, according to the manufacturer’s instructions, to immunoprecipitate proteins of interest with either mouse anti human FN1, AR or cMYC monoclonal antibody or IgG (negative control). Immunoprecipitated proteins was cleaved from the crosslinker with DTT at 50mM in SDS-PAGE sample buffer. Proteins immunoprecipitated were analyzed with Western Blot technique as described by Das *et al*., using rabbit anti human polyclonal FNDC1 (PA5-56603, Invitrogen, UK), FN1 (ab2413, Abcam, USA), AR and cMYC (10828-1-AP, Proteintech, USA) antibody.

### Transient Knockdown of FN1, AR, cMYC and Overexpression of miR-1207-3p

The castration resistant C42B cells harvested in complete media were transiently transfected using siRNA oligonucleotides, 25nM in Opti-MEM using Lipofectamin RNAimax (Invitrogen, USA) transfection reagent for 48-72 hours. Knockdown of FN1, AR, cMYC was performed using their respective siRNA oligonucleotides (L-003282-02-0010, Horizon Discovery Biosciences Limited, UK). In addition to that silencing with non-targeting pool of siRNA oligonucleotides (D-001810-10-20, Horizon Discovery Biosciences Limited, UK) and GAPDH siRNA oligonucleotides (D-001830-10-20, Horizon Discovery Biosciences Limited, UK) served as negative and positive control respectively for any observed knockdown. Levels of FN1, AR, cMYC knockdown were analyzed by immunoblotting using respective rabbit anti human primary antibodies. To analyze the effect of FN1, AR, cMYC knockdown on FNDC1/FN1/AR/cMYC interaction, siRNA transfected cells were subjected to co-immunoprecipitation assay.

Cells were seeded in 150mm plates until 30% - 40% confluency is reached. Cells were transfected with Lipofectamin RNAimax either a 50nM non-targeting negative control oligonucleotide (MISSION® Synthetic microRNA Negative Control, product# NCSTUD001) or 50nM miR-1207-3p oligonucleotide mimic (MISSION® microRNA Mimic, product# HMI0066) for 48-72 hrs in Opti-MEM (Thermo Fisher Scientific Inc; Wilmington, DE, U.S.A) combined with complete media.

### Immunofluorescence

Glass cover slips sterilized with 70% ethanol were used to seed cells. Cells were allowed to adhere for 2 days and fixed with 4% formaldehyde for 1 hour followed by washing 3 times for 5 minutes. Next the fixed cells were permeabilized with .5% Tritonx-100 followed by washing. Permeabilized cells were then blocked for an hour in Abcam blocking buffer. Thereafter cells were incubated with primary antibody in 3% (w/v) BSA/PBS (1:100), washed with 1% (w/v) BSA/PBS, and subsequently incubated with species specific fluorescently tagged secondary antibody in 1% (w/v) BSA/PBS (1:500). Primary antibodies used are same as the western blot experiments. Nuclei were stained with Vectashield Plus Antifade Mounting Medium with DAPI (H-2000). Images were captured using the Nikon eclipse Ti confocal microscope and analyzed using NIS-Elements Confocal. Secondary antibodies used in fluorescent staining were Alexa Fluor Plus 488 (A32731), Alexa Fluor Plus 647 (A32733) and Alexa Fluor Plus 555 (A32732), all against Rabbit.

### Ethics statement

This study does not include any research involving any human participants. Consequently, there was no need for consent from anyone, and no minors were studied. Commercially obtained cell lines were used for this study.

## Results

### Interaction Between FNDC1/FN1/AR/cMYC In Prostate Cancer Cells

Previous study in our lab has shown that inhibition of FNDC1 decreases endogenous FN1, AR and cMYC expression. Thus, to determine if there is any interaction between FN1/FNDC1/AR/cMYC we performed a co immunoprecipitation (**Figures 1A, B and C**). Cells fixed with BS3 were subjected to lysis and the protein extracts obtained from the non-tumorigenic prostate epithelial RWPE1 and prostate cancer cells LNCaP (metastatic prostate cancer cell line) and C4-2B (the castration resistant cell line) were incubated with monoclonal anti-FN1, anti-AR and anti-cMYC antibody coupled Dynabeads. Western Blot analyses of the immunoprecipitated proteins for anti FN1 resulted in single bands at 285KDa (FN1), 265KDa (FNDC1), 110KDa (AR-B) and 55KDa (cMYC). For anti-cMYC at 285KDa, 265KDa, 72KDa (AR-A) and 55KDa and that for anti-AR the bands observed are at 285KDa, 265KDa, 110KDa, 72KDa and 55KDa in LNCaP and C42B. Thus, we can surmise that FN1/FNDC1/AR/cMYC interacts in prostate cancer cell lines. However, only FN1/AR/cMYC interacts in the normal prostate cell line. Non-immunoprecipitated cell lysates served as input and incubation of IgG coupled Dynabeads with cell lysates acted as the negative control. It is interesting to note that in the PCa cell lines, FN1 interacts only with full length AR-B and cMYC interacts only with spliced AR-A which is phenotypical of prostate cancer cells.

**Figure 1.**
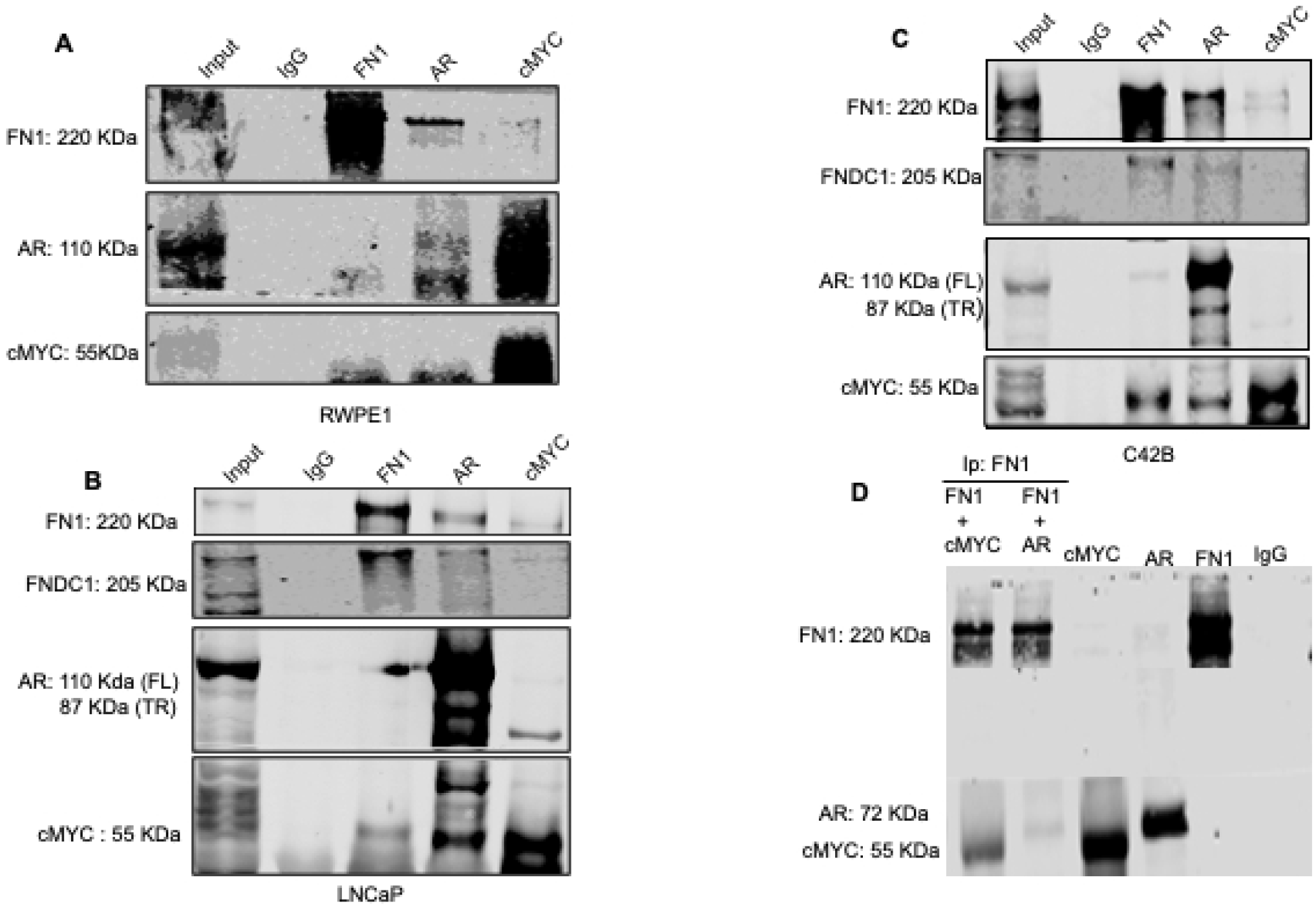
Interaction between FNDC1, FN1, AR, and cMYC in RWPE1, LNCaP and C42B. **(A), (B) & (C)** Cells from RWPE1 normal prostate epithelial cell line and PCa cell lines were subjected to lysis and then pulled down with IgG, FN1, AR and cMYC mouse monoclonal antibodies as indicated across the horizontal axis. Whereas the vertical axis represents proteins that were probed with specific antibodies with the help of western blot. FNDC1/FN1/AR/cMYC interacts with another. **(D)** Anti FN1 or IgG coupled Dynabeads were incubated with FN1, AR and cMYC. Co-immunoprecipitated samples resolved via immunoblotting. FN1, AR and cMYC were loaded as input controls.

To further corroborate our observations, we performed an *in vitro* coimmunoprecipitation (**Figure 1D**) with recombinant proteins. The purified proteins (FN1, AR, cMYC) served as the input and IgG as negative control. Incubation of anti-FN1 coupled Dynabeads with recombinant proteins FN1 and AR resulted in a single band for each. Similarly, a single band for FN1 and cMYC showed up in the second set of the same experiment with FN1 and cMYC recombinant proteins. This further confirmed FN1/AR/cMYC interaction.

### Direct physical interaction between FNDC1/ FN1 /AR/cMYC

Next, we wanted to investigate the effect of transiently silenced FN1 on the FN1/FNDC1/AR/cMYC interaction in C4-2B cell. Cells transfected with non-target siRNA oligonucleotides served as negative control whereas transfection with GAPDH served as positive control to confirm successful knockdown. Cell lysates from negative, positive and FN1 knockdowns respectively were subjected to co-immunoprecipitation with monoclonal FN1 antibody and the untreated was included as input. In addition, cell lysates from FN1 knockdown cells were immunoprecipitated with monoclonal AR and cMYC antibodies. Analyses of the protein by immunoblot using polyclonal FN1, FNDC1, AR and cMYC antibodies respectively revealed FN1/FNDC1/AR/cMYC interaction in positive and negative controls but not in the FN1 knockdown (**Figure 2**). In case of AR pull down a faint interaction between FN1 and AR and normal interaction between AR and cMYC was observed. Pull down with cMYC showed interaction only between AR and cMYC. This led us to believe that FNDC1 has direct physical interaction only with FN1, whereas there is a direct interaction between FN1/AR/cMYC. We did similar experiments with AR and cMYC knockdowns as well (**Supplemental figure 2**).

**Figure 2.**
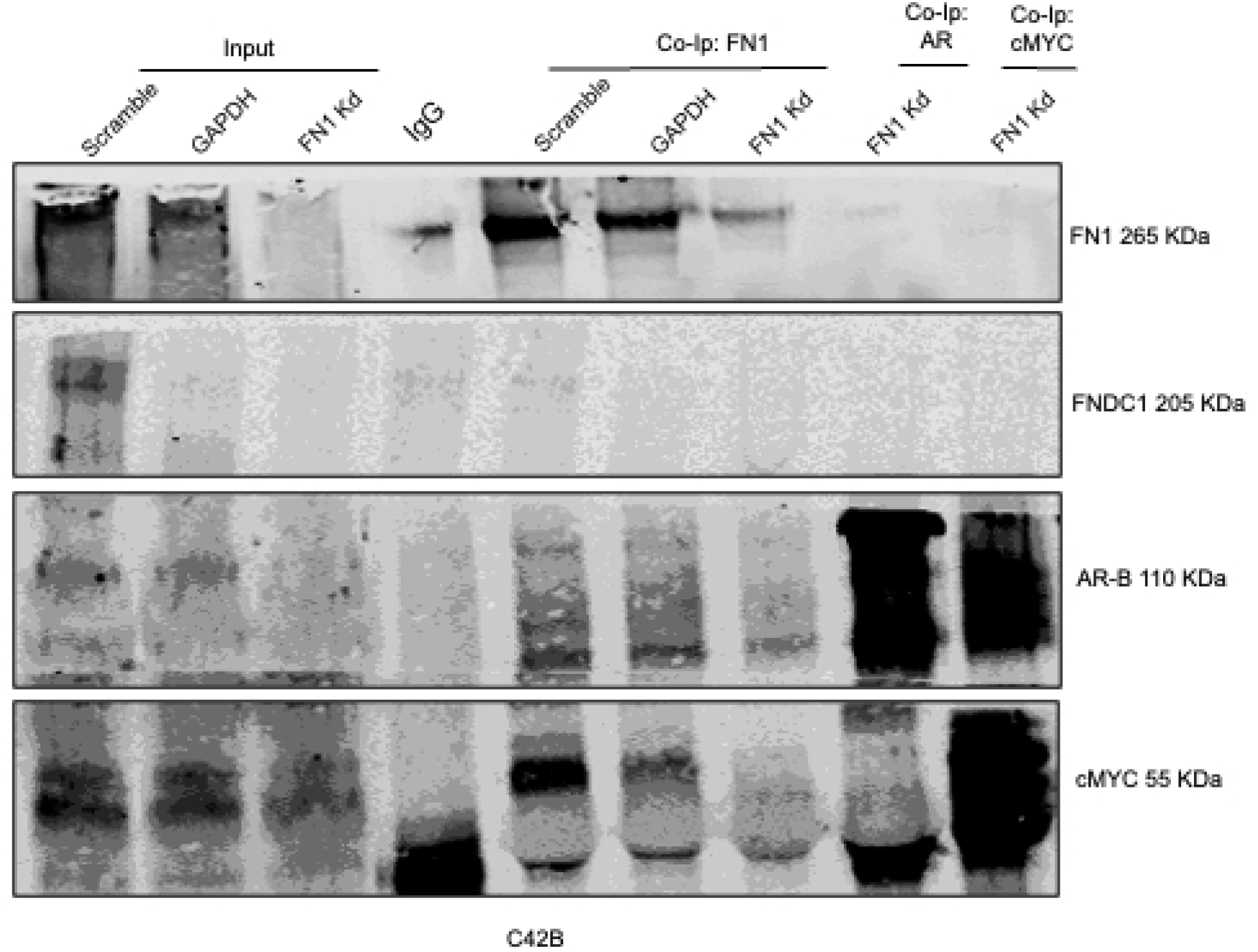
Effect of knockdown on FNDC1, FN1, AR, and cMYC interaction in C42B. C42B cells were transfected with a pool of siRNA against FN1. GAPDH acted as positive control and scramble non-targeting as a negative control. After 72 hours cells were subjected to lysis and then pulled down with IgG, FN1, AR and cMYC mouse monoclonal antibodies. Pulled-down samples were analyzed by immunoblotting with rabbit anti human FNDC1, FN1, AR and cMYC. FNDC1/FN1/AR/cMYC has direct physical interaction between one another.

### Intracellular localization of FNDC1/FN1/AR/cMYC in prostate cancer cells

Following the confirmation of the interaction between FNDC1/FN1/AR/cMYC we wanted to investigate the intracellular localization of these proteins in normal prostate epithelial (RWPE1), non-CRPC (LNCaP) and CRPC (C4-2B) cells. FNDC1, FN1, AR and cMYC expression was detected in the normal as well as the two PCa cell lines (**Figure 3**). The RWPE1 cell line showed lower levels of all the four proteins compared to the PCa cell lines as confirmed by previous immunoblotting data. FN1 is observed in the cytosol of all three cell lines. A dispersed intracellular FN1 localization pattern was observed in the LNCaP cells whereas in RWPE1 and C42B cells it is observed to be perinuclear. AR and cMYC have been shown to reside both in the cytosol and the nucleus which was confirmed via the dispersed staining pattern as seen for AR and cMYC in all three cell lines. FNDC1 showed a distinct diffused localization pattern in the cancer cells distinct from its nuclear localization in the non-tumorigenic cell line. Thus, the data led us to believe that the interaction between the FNDC1/FN1/AR/cMYC is plausible in the cytosol of the PCa cell line.

**Figure 3.**
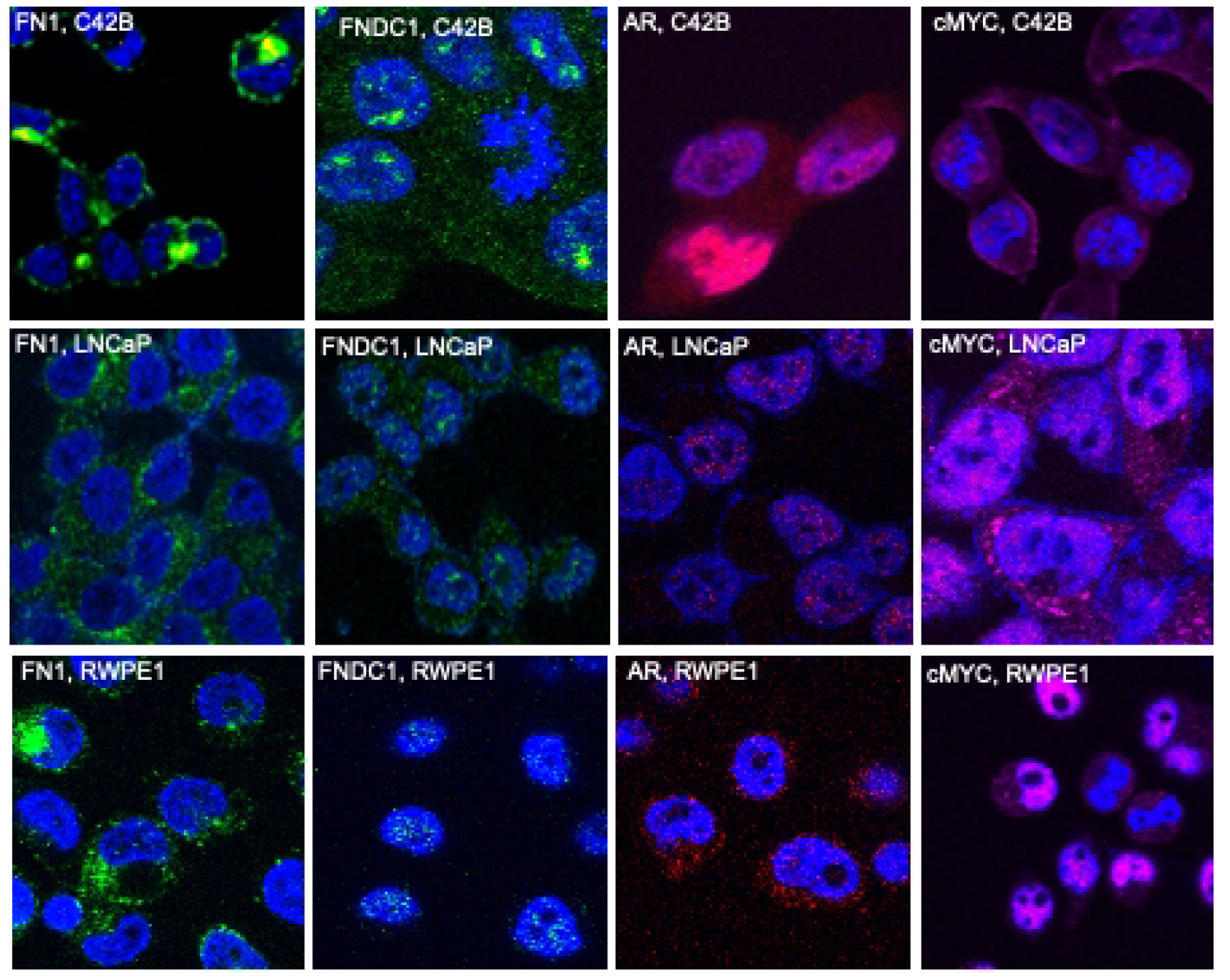
Localization of FNDC1, FN1, AR, and cMYC in RWPE1, LNCaP and C42B. RWPE1 normal prostate epithelial cells and LNCaP and C42B PCa cells were stained with specific antibodies for FNDC1, FN1, AR, and cMYC. FN1 and FNDC1 are indicated by green, cMYC by magenta and AR by red. FNDC1, AR and cMYC are observed in the nucleus as well as in the cytoplasm in the PCa cells. Interestingly FNDC1 is only observed in the nucleus in RWPE1. FN1 is observed only in the cytoplasm in all three cell lines. Scale bars are equivalent to 10µm.

### The Effect of MiR-1207-3p Overexpression on FN1/FNDC1/AR/cMYC Interaction

Collectively the interaction data, the intracellular localization and FNDC1 being a direct molecular target of miR-1207-3p led us to hypothesize that miR-1207-3p might have an effect on the interaction between FNDC1/FN1/AR/cMYC. To validate our hypothesis C42B cells were subjected to miR1207-3p overexpression via exogenous introduction of mRNA mimic in the cell. MiR-1207-3p overexpression was confirmed by qRT-PCR which showed 10-fold increase in the miRNA in the mimic transfected compared to non-scramble one. Monoclonal AR antibody was used to pull down FNDC1/FN1/AR/cMYC proteins in both NC and mimic transfected cells (**Figure 4**). The immunoprecipitation data didn’t reveal any significant change in the interaction between F(f)AM complex. Thus, we can conclude that miR-1207-3p does not have any effect on the FNDC1/FN1/AR/cMYC interaction.

**Figure 4.**
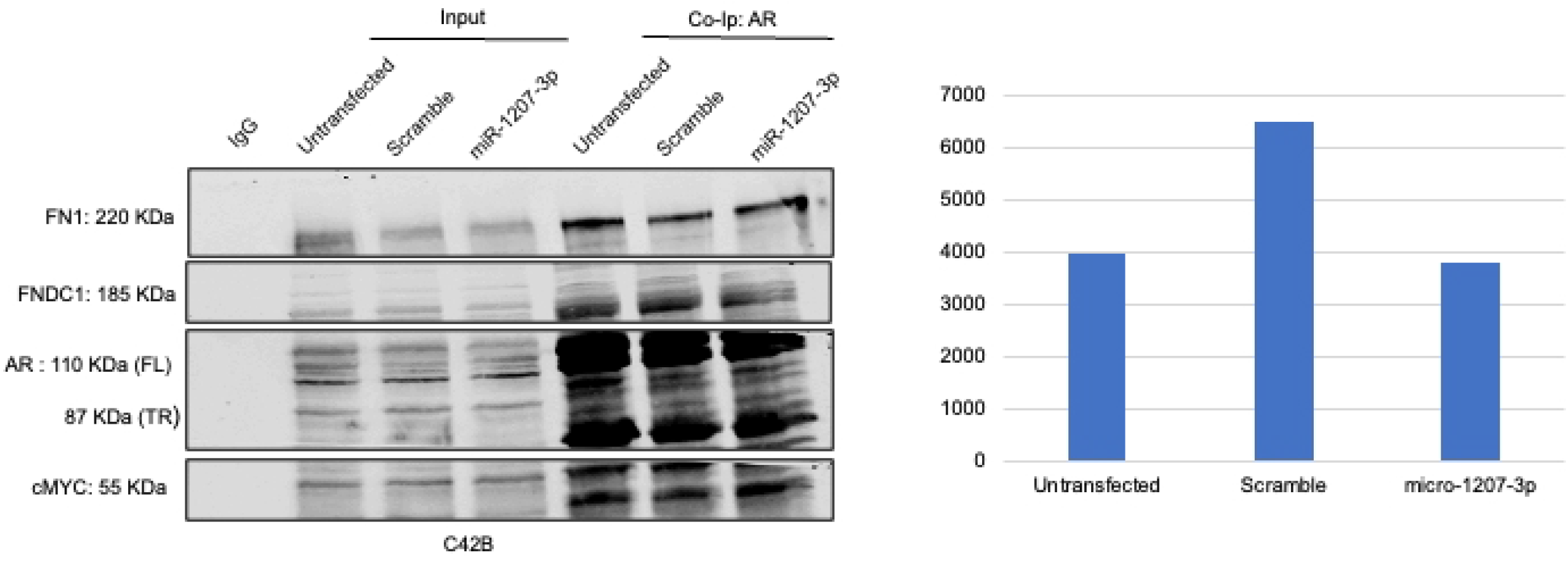
Effect of miR1207-3p overexpression on FNDC1, FN1, AR, and cMYC interaction in C42B. Adherent C42B cells were treated with miR-1207-3p mimic. Cells treated with miR-1207-3p, scramble and untreated cells were subjected to lysis and then pulled down with IgG and AR mouse monoclonal antibodies as indicated across the horizontal axis. Whereas the vertical axis represents proteins that were probed with specific antibodies via western blot. The graph plot shows reduction in FNDC1 due to exogenous overexpression of miR-1207-3p. miR-1207-3p overexpression affects FNDC1 expression but not FNDC1/FN1/AR/cMYC interaction.

## Discussion

Fibronectin is a well-studied protein molecule that mediates cellular interaction with the ECM and AR/cMYC are well known transcription factors. On the other hand, more studies are being conducted to understand the role of FNDC1 in cells. Ours is the first study to demonstrate intracellular interaction between FN1 FNDC1, AR and cMYC in prostate cells. The capacity of FN1 to promote cellular activities relies on its ability to interact with different macromolecules [8]. Fibronectin can act as a ligand to a variety of molecules including integrins [14], growth factor receptors [15,16]. In addition to cell receptors FN1 is also known to interact with different ECM components such as gelatin, collagen, fibrillin etc. [9,10,13]. cMYC a well-studied transcription factor is known to take part in several interactions that are transcription independent. Such as the promotion of mRNA cap methylation, stimulation of DNA replication, interaction with tumor suppressors or cell cycle regulators and ubiquitin ligases [18,19,20]. Therefore, a study was conducted to look at the different interactors of cMYC. Among the various cMYC interactors identified in human fibroblast cells with the help of SILAC technique, FN1 was revealed as one of the interactors [17]. Co-immunoprecipitation with FN1, AR and cMYC antibodies suggested an interaction between FNDC1, FN1 and AR and cMYC. In addition to that, *in vitro* coimmunoprecipitation with recombinant proteins further confirmed the interaction between FN1, AR and cMYC, although FNDC1 could not be investigated because recombinant FNDC1 is unavailable.

cMYC interacts with AR-A, the truncated isoform but not with full length AR. Concordant with our data, Bai *et al*. was able to show that cMYC enhances AR-FL and AR-V protein stability and regulates AR expression but they could not show a direct physical interaction between AR-FL/AR-V and cMYC [24]. Fibronectin increased cMYC expression in BEA-2B human bronchial epithelial cells [22]. Therefore, we conducted a knockdown of FN1, AR and cMYC which not only confirmed interaction between FN1/AR/cMYC, but also showed that there is direct physical interaction between FN1/AR/cMYC. We were unable to fully establish the direct physical interaction of FNDC1 with FN1, AR and cMYC.

Scientists reported cMYC translocation in B-cell lymphoma when they observed primarily nuclear but also mixed nuclear and cytoplasmic staining pattern of cMYC in these cells [2]. Ligand binding is necessary for AR to move to the nucleus where it can access the androgen responsive gene and regulate them [3], [4], [5]. AR is exported back to the cytoplasm at the withdrawal of ligand [6]. Presence of extracellular, intracellular and membrane bound Fibronectin was observed in glomerular cells [7]. Taken together, the above-mentioned studies validate our findings, FN1 localizes to the cytoplasm of both non-tumorigenic as well as tumorigenic prostate cells, whereas AR as well as cMYC is observed in the cytoplasm and nucleus. Of importance is the localization of FNDC1, which is only observed in the nucleus of non-tumorigenic cell line RWPE1 but both nucleus and cytoplasm of the prostate cancer cells. Collectively, this data show that FNDC1/FN1/AR/cMYC.

Multiple studies have shown that miRNAs are capable of modulating signaling cascades by targeting 3’UTR of one of the signaling molecule in a particular signaling pathway. Additionally, Dibash *et al*. had shown that FNDC1 is a direct molecular target of miR-1207-3p. Moreover, overexpression of miR-1207-3p reduces FNDC1, FN1, AR and cMYC expression whereas inhibition increases the expression of these proteins[31]. Therefore, we were interested to study the effect of miR-1207-3p on the FNDC1/FN1/AR/cMYC. Although miR-1207-3p overexpression has no direct effect on FN1/FNDC1/AR/cMYC interaction yet it decreased FNDC1 expression. This, could be because FN1, AR, cMYC does not have a strong affinity for FNDC1.

The mechanism by which FNDC1/FN1/AR/cMYC interacts is currently uncharacterized. The NTD domain of the AR receptor is reported to be important in the activation of the LBD. Apart from the AF1, which is the main transcription activation domain of AR, there are two other units Tau-1 and Tau-5. Our initial interest was to see if FN1 can act as a ligand to AR. However, our results show that FN1 can only interact AR-B/AR-FL but not to the truncated AR-A. This led us to believe that FN1 associates itself with AR through the NTD of AR [26]. Fibronectin interacts with integrin receptors through the central binding domain (FNIII_1-12_) [27, 28]. On the other hand, HSP90 a chaperone protein interacts with FN1 via the N-terminal regions to aid in matrix assembly [29]. Thus, we can predict that FN1 can interact with AR through the CBD of FN1. The above-mentioned studies also help us to predict that it is possible FN1 interacts with cMYC by its N-terminal domain and AR through any domain other than the N-terminal domain. It was not easy to determine which domain of cMYC is possibly involved in the interaction with FN1 and AR as multiple domains of cMYC are involved in similar functions [30]. It will be interesting to confirm the domains of the three proteins that are involved in the interaction through mutational studies. We already know that two different isoforms of AR are involved in the F(f)AM complex. A study in *Drosophilla* revealed that cell growth and apoptosis is differentially regulated by the different human cMYC isoforms [32]. Hence, one intriguing possibility is that different isoforms of cMYC are involved in the FNDC1/FN1/AR/cMYC. It is interesting to note that cMYC in the cytoplasm is known both to induce apoptosis via cytochrome as well as sensitize cells to apoptosis [25]. Therefore, we won’t be completely wrong to assume that cMYC present in the cytoplasm of the prostate cells is possibly involved in apoptosis.

In conclusion, FNDC1/FN1/AR/cMYC can be important in mediating cellular function in cells, further study is required to establish the role of this interaction in prostate epithelial cells. Moreover, we can surmise that there is a possibility of two separate pathways by which FNDC1/FN1/AR/cMYC regulates cellular functions (**Figure 5**), one in which FN1 interacts with FNDC1, AR-B (AR full length) and cMYC and the other will be c-MYC interaction with FN1 and AR-A (AR truncated).

**Figure 5.**
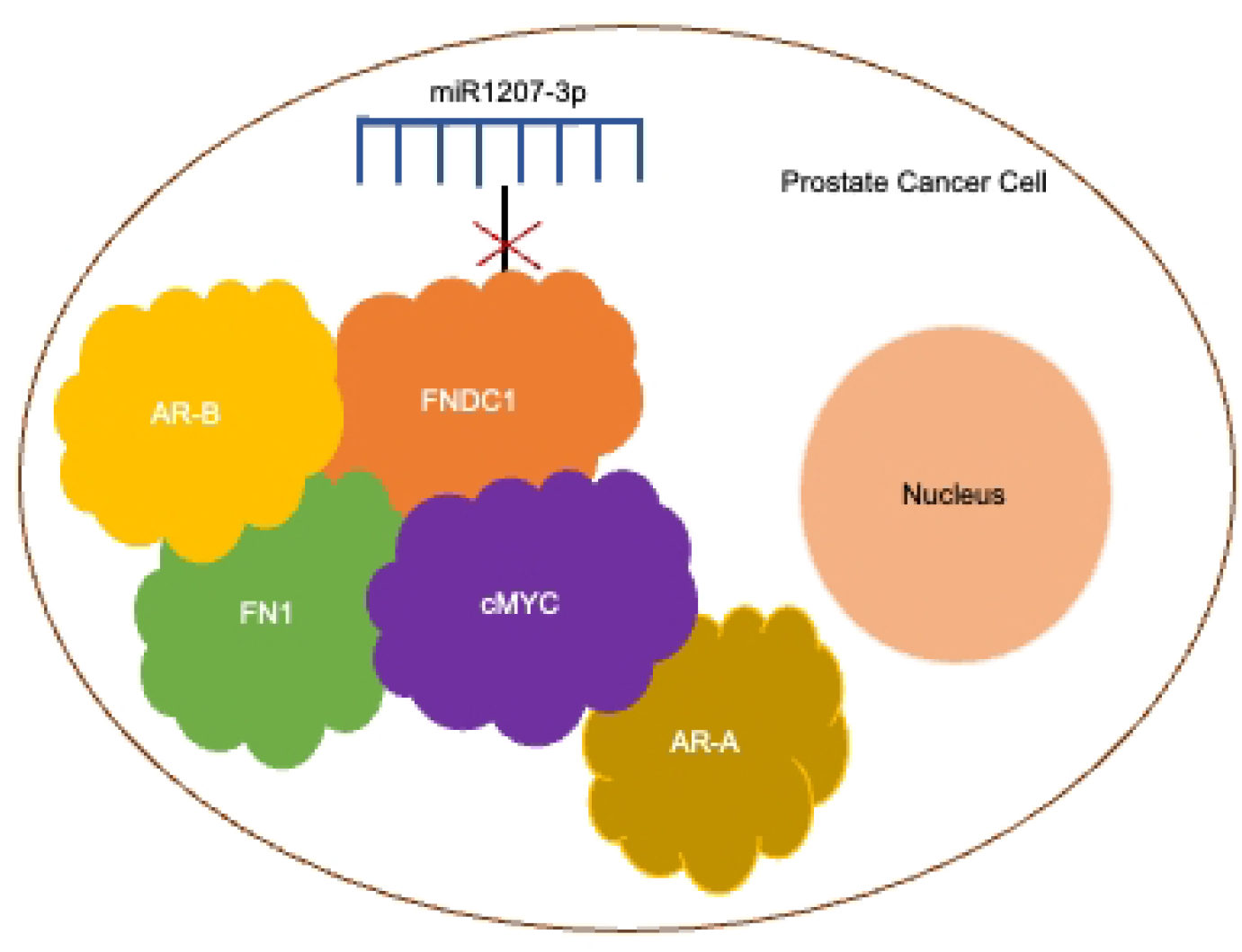
The interaction between FNDC1, FN1, AR and cMYC is not regulated by miR-1207-3p. The cartoon depicts the possibility of two different pathways. One in which FN1 interacts with FNDC1, AR-B (AR full length) and cMYC and the other in which c-MYC interacts with FN1 and AR-A (AR truncated).

